# Patterns of connectome variability in autism across five functional activation tasks. Findings from the LEAP project

**DOI:** 10.1101/2022.02.22.481408

**Authors:** Tristan Looden, Dorothea L. Floris, Alberto Llera, Roselyne J. Chauvin, Tony Charman, Tobias Banaschewski, Declan Murphy, Andre. F. Marquand, Jan K. Buitelaar, Christian F. Beckmann, the AIMS-2-TRIALS group

## Abstract

**Background:** Autism spectrum disorder (autism) is a complex neurodevelopmental condition with pronounced behavioural, cognitive, and neural heterogeneities across individuals. Here, our goal was to characterise heterogeneity in autism by identifying patterns of neural diversity as reflected in BOLD fMRI in the way individuals with autism engage with a varied array of cognitive tasks.

**Methods:** All analyses were based on the EU-AIMS/AIMS-2-TRIALS multisite Longitudinal European Autism Project (LEAP) with participants with autism and typically developing controls (TD) between 6 and 30 years of age. We employed a novel task-potency approach which combines the unique aspects of both resting-state fMRI and task-fMRI to quantify task-induced variations in the functional connectome. Normative modelling was used to map atypicality of features on an individual basis with respect to their distribution in neurotypical control participants. We applied robust out-of-sample canonical correlation analysis (CCA) to relate connectome data to behavioural data.

**Results:** Deviation from the normative ranges of global functional connectivity was greater for individuals with autism compared to TD in each fMRI task paradigm (all tasks p<0.001). The similarity across individuals of the deviation pattern was significantly increased in autistic relative to TD individuals (p<0.002). The CCA identified significant and robust brain-behavior covariation between functional connectivity atypicality and autism-related behavioral features.

**Conclusions:** Individuals with autism engage with tasks in a globally atypical way, but the particular spatial pattern of this atypicality is nevertheless similar across tasks. Atypicalities in the tasks originate mostly from prefrontal cortex and default mode network regions, but also speech and auditory networks. We show, moving forward, sophisticated modeling methods such as task-potency and normative modeling will prove key to unravelling complex heterogeneous conditions like autism.

## Introduction

Autism spectrum disorder (henceforth ‘autism’) is a complex neurodevelopmental condition marked by difficulties with social communication, repetitive, restricted behaviours and interests and sensory processing atypicalities (American Psychiatric Association, 2013). Cross-participant heterogeneity in autism has made understanding underlying mechanisms and the complex interrelation between neurobiology and cognitive profiles in autism challenging. Imaging studies in autism report both over-and under-connectivity of functional brain networks (Oldehinkel et al., 2019; Picci et al., 2016; Uddin et al., 2013) on the basis of resting-state fMRI data. Different task-fMRI studies, probing a variety of neural processes, report between-group differences with small effect size at best (Deshpande et al., 2013; Just et al., 2014; Moessnang et al., 2020). Crucially, little effort has been made to integrate the diverse findings both across different cognitive domains and between task-fMRI and resting-state connectivity at the level of an individual participant. In order to better characterize heterogeneity both across cognitive domains and across individuals we combine novel methodological approaches.

First, we propose an integrated analytical approach to characterise the task-specific cognitive demands in autistic individuals. We utilise a task-potency approach which combines the unique aspects of both resting-state fMRI (rs-fMRI) and task-fMRI (Chauvin et al., 2019). Rs-fMRI provides insight into the large scale ‘architecture’ of brain connectivity in an individual. Task-fMRI might however more directly probe the neural correlates of specific cognitive domains affected by the condition such as social/emotional processing and attention. We leverage the advantages offered by both views in task-potency, which disentangles the relative contribution of task-induced functional connectivity from that of the baseline architecture at the individual level (Mennes et al., 2010). This allows for greater sensitivity to individual task-based functional connectivity (FC) effects as well as a more precise interpretation of findings as being related specifically to the cognitive load and not to differences in baseline.

Second, even though many cognitive/behavioural studies have been able to successfully show differences between individuals with autism and typically developing individuals across a range of cognitive domains such as social cognition, reward and emotion processing, and executive functioning (Hull et al., 2017), there appears to exist pronounced behavioural, cognitive, and neural heterogeneity across individuals with autism (Brunsdon & Happé, 2014; Nunes et al., 2018; Wolfers et al., 2019). In order to parse the heterogeneous nature of autism neurobiology, we therefore apply normative modeling which will allow us to map atypicality of brain-derived features on an individual basis with respect to the distribution of those features in a group of similar typically developing controls. (Marquand et al., 2019). This approach has previously been applied in autism and yielded promising results (Bethlehem et al., 2020; Floris et al., 2020; Zabihi et al., 2019). This way, the analysis becomes more sensitive to idiosyncratic brain atypicalities.

In order to be able to characterise diversity in presentation across cognitive domains and individuals we leverage the large-scale resource that has specifically been designed to capture a large, heterogenous and thus naturalistic autism sample – the EU-AIMS/AIMS-2-TRIALS Longitudinal European Autism Project (LEAP) (Charman et al., 2017a, 2017b; Loth et al., 2017). It is designed to provide deep-phenotyping by including both various neuroimaging measures (such as rs-fMRI and task-fMRI), an extensive cognitive battery capturing social cognition, reward and emotion processing, and executive functioning and in-depth behavioural phenotyping. Due to the presence of multiple task paradigms in the dataset we are able to contrast the spatial patterns of atypicality across these different tasks. We are especially interested in finding out whether posited patterns of task-specific functional connectome atypicality in autism are similar across cognitive domains – and conversely, how that similarity is expressed in typically developing controls.

We assess whether the patterns of brain atypicality we find in individuals relate to metrics of autism at a behavioural level - thereby assessing for each task whether task-specific functional connectome atypicality carries information relevant to finding brain-behaviour relationships in autism. In order to relate high-dimensional brain data to behavioural data in a way without making prior assumptions on the most relevant features in a multivariate context, we will apply canonical correlation analysis (CCA) (Mihalik et al., 2020; Wang et al., 2020)

By integrating complementary functional modalities and combining the aforementioned novel tools such as task-potency and normative modelling, taking a unique look at identifying (a)typicality in the way individuals with autism engage with cognitive demands across tasks at the individual neural level is made possible. For the multiple fMRI tasks present in the dataset enable a crucial cross-task perspective towards gauging to what extent such atypicalities exist across different cognitive loads, and whether this pattern is different in autism from typically developing controls.

## Materials and Methods

### Sample

The dataset from the EU-AIMS/AIMS2TRIALS LEAP project was used for the current analyses – a large multi-centre European project with an aim to identify biomarkers in (Charman et al., 2017b, 2017a; Loth et al., 2017). Local ethical committees approved the study at their respective sites. Participant were extensively clinically phenotyped and underwent multiple MRI assessments. Data from participants with intellectual disability (IQ<75) was not included in the current project. Furthermore, we removed participants from the analysis on the basis of data quality using the following criteria: Participants were required to have acceptable overlap (>94%) with the MNI152 standard brain after image registration. We then excluded 57 participants due to poor overlap (<50%) with one or more particular regions from the ICP brain parcellation atlas. Participants were furthermore excluded on the basis of incidental findings and incomplete scans (N=18), and those in the top 5% in terms of head motion quantified through root mean square framewise displacement (N=27) (Jenkinson et al., 2002). The above criteria resulted in the inclusion of data for analyses from the following participants: 282 participants with autism (age range 7.5-30.3 years; mean = 17.1; sd = 5.4; 72.3% male), and 221 typically developing controls (age range 6.9-29.8 years; mean = 17.0; sd = 5.5; 63.8% male). Not every participant was included in each analysis, for the reason that not all participants completed all fMRI scan sessions. See supplementary tables S1 through S5 for more details on the subsample characterization in each task.

### Behavioural data

Total scores on seven behavioural variables were included in the multivariate canonical correlation analyses with the aim to broadly include information about the affected autism domains, co-occurring attention deficit hyperactivity disorder (ADHD), and adaptive functioning, as well as IQ. Three variables cover the primary affected domains in autism (social and communicative difficulties, repetitive/restricted behaviours and interests, and sensory atypicalities): Social Responsiveness Scale-2 (SRS) (Constantino, 2013) – a quantitative scale of autism symptomatology over the past 6 months, the Repetitive Behaviors Scale-revised (RBS) (Bodfish et al., 2000) – a scale which assesses more specifically restricted and repetitive behaviours, and the Short Sensory Profile (SSP) (Tomchek & Dunn, 2007) – a scale which assesses sensory processing atypicalities. Two further variables cover the ADHD related behaviours: ADHD hyperactivity/impulsivity, and ADHD inattentiveness. These are the two components from the DSM-5 ADHD rating scale for behaviour in the past six months. Furthermore, we included the Vineland-II Adaptive Behaviour Composite (Vineland) (Sparrow, 2011) which assesses the level of real-life everyday adaptive functioning. Finally, we included Full scale intelligence quotient (IQ) as measured by the Wechsler adult scale intelligence-2 (Wechsler et al., 2011). Missing data was imputed with random forest regression, as done in (Zabihi et al., 2019).

### fMRI data

Participants performed a resting-state fMRI (rsfMRI) lasting approximately 6 minutes, and one or more of the following task-fMRI scans: Hariri emotion processing (Hariri) (Hariri et al., 2002), Flanker and Go-NoGo (Flanker) (Meyer-Lindenberg & Weinberger, 2006), social reward anticipation (Reward_s), non-social reward anticipation (Reward_ns) (Delmonte et al., 2012), and animated shapes theory of mind (ToM) (Castelli et al., 2002; White et al., 2011). See supplementary information for brief descriptions of these tasks. Participants were instructed to relax and fixate on a cross presented on the screen for the duration of the rsfMRI scan. Additionally, each participant completed an anatomical scan for the purpose of registration. MRI data were acquired on 3T scanners at multiple sites in Europe - King’s College London (KCL), Radboud University Nijmegen Medical Centre (RUNMC), University Medical Centre Utrecht (UMCU), Autism Research Centre (ARC), University of Cambridge (UCAM), Central Institute of Mental Health, Mannheim (CIMH), and Karolinska Institutet (KI). Data from KI was not used due to a low number of participants. fMRI parameters are described in the supplementary information.

### fMRI preprocessing

Preprocessing of both the resting state- and task-fMRI data was performed with tools from FSL (Jenkinson et al., 2012). The first five volumes for each acquisition were removed to allow for equilibration of the magnetization. To correct for head movement, we performed volume realignment to the middle volume using MCFLIRT. Next, global 4D mean intensity normalization and smoothing with a 6mm FWHM kernel were applied. ICA-AROMA was used to identify and remove secondary motion-related artefacts (Pruim, Mennes, Buitelaar, et al., 2015; Pruim, Mennes, van Rooij, et al., 2015). Next, signal from white matter and cerebrospinal fluid was regressed out and we applied a 0.01Hz high-pass filter. For each participant, we registered acquisitions to their respective high-resolution T1 anatomical images by means of the Boundary-Based Registration tool from FSL-FLIRT (Jenkinson et al., 2002). The high-resolution T1 image belonging to each participant was registered to MNI152 space with FLIRT 12-degrees of freedom linear registration, and further refined using FNIRT non-linear registration (Andersson et al., 2007). We used the inverse of these transformations to take a brain atlas to the native space of each participant, where all further analyses were performed. ComBat was used to clean the data for linear site effects (Johnson et al., 2007)

### Task potency

We used the hierarchical instantaneous connectivity parcellation (ICP) brain atlas with 168 brain regions (hierarchically situated in 11 larger-scale ‘networks’) (van Oort et al., 2017) to define brain regions in the native space of each subject. This was done for each of the five task-fMRI acquisitions – as well as for resting-state. For each participant, we calculated the regularized covariance (Ledoit & Wolf, 2012) between the average BOLD time series extracted from each brain region pair. We then estimated partial correlations from the covariance matrix and consecutively applied the Fisher-Z transformation. This provided for each participant a connectivity matrix of size 168×168 - one resting-state matrix and one task-fMRI matrix per task. The main gaussian from a mixture gaussian-gamma model that was fitted on the distribution of edge values for each individual matrix supplied us with parameterised information about said distribution (Bielczyk et al., 2017; Llera et al., 2016). We used these parameters to normalize the elements in each matrix. In order to produce individual matrices of connectivity modulations induced by the task, i.e., task potency, we subtracted each participant’s resting-state connectivity matrix from that participant’s task connectivity matrix. The resulting matrices are interpreted as containing the connectivity modulations away from the resting-state baseline that the respective task induces in the brain – i.e. task-potency (Chauvin et al., 2019, 2021).

### Normative modelling

A normative model of potency edge values was built against age and sex from typically developing participants using the nispat implementation in python (Marquand et al., 2016, 2019). This model was used to predict the range of (a)typical edge values in out-of-sample typically developing participants as well as the autistic participants. Task potency values were thereby transformed to Z-scores quantifying the atypicality per edge of each individual’s task potency matrix elements given the normative reference. This was done separately in each task. Percentage of connectome deemed atypical was identified by thresholding the Z-scores at z=|±2.571|, nominally describing the 1% most extreme positive and negative values. The number of edges passing this threshold for each individual was then expressed as a percentage of the total amount of edges. Independent t-tests were performed on the percentages to assess case-control differences in the tasks. All p-values were FDR corrected for multiple comparisons across tasks. Cohen’s D was calculated for an effect size estimate in each task.

### Cross-task similarity

We investigate similarity in the patterns of atypicality across the tasks in autism as well as typically developing controls. This is done for both diagnostic groups separately by constructing a pearson correlation matrix from the mean edge-wise atypicality scores across tasks. Differences in covariance values are compared between the groups with the Wilcoxon signed rank test (Wilcoxon, 1945).

### Canonical correlation analysis

Canonical correlation analysis (CCA) is a technique for finding latent linear multivariate relationships between two sets of data (Hotelling, 1936; Pedregosa et al., 2011; Smith et al., 2015). CCA analysis was done only in the individuals with autism in order to assess whether the variation in the task atypicality patterns relates also to phenotypic description in individuals with autism. The following processing steps were done separately for each task. For the behavioural side of the CCA analysis the seven variables previously described were used (SRS, RBS, SSP, ADHD hyperactivity/impulsivity, ADHD inattentiveness, Vineland, IQ). For the brain side of the CCA, principal components-based dimensionality reduction was applied to the brain subjects by edges data matrix in order to make the CCA well-posed. Aiming for a balance between model accuracy and complexity, the top 10 variance components were kept for input in the CCA. Stability and generalizability of CCA parameters was assessed through 1000 different splits of 10-fold out-of-sample cross validation – building a ‘test’ distribution of the first canonical correlation as well as the paired CCA weights. Significance of the relationship found was assessed through building an out-of-sample null distribution of CCA correlations i.e., permutation testing. Subjects were randomly permuted (within scan site) to break up the original correlation structure and performing again 1000 different splits of 10-fold cross-validation. Non-spuriousness of the original relationship found was assessed by assessing whether the mean of the test distribution is more extreme than the 95th percentile of the null distribution.

## Results

### General connectome atypicality levels

Figure 1 shows for each task distributions of subjects for the percentage of total edges that pass the atypicality threshold at z=2.571 (1%, two tailed) in the autism and TD groups. We interpret this metric as a global subject-level atypicality score in each task. We identify significantly greater levels of atypical modulation in individuals with autism in each of the tasks. Flanker - Cohen’s d: 0.61, p<0.01. Hariri - d: 0.34, p<0.01. Monetary reward - d: 0.19, p<0.01. Social reward - d: 0.19, p<0.01. Theory of mind - d: 0.36, p<0.01. For full information see supplementary table T1. These findings provide a global view where the distribution of edges in autism is shifted towards greater atypicality in achieving the same cognitive states as represented by task-potency.

**Figure 1.**
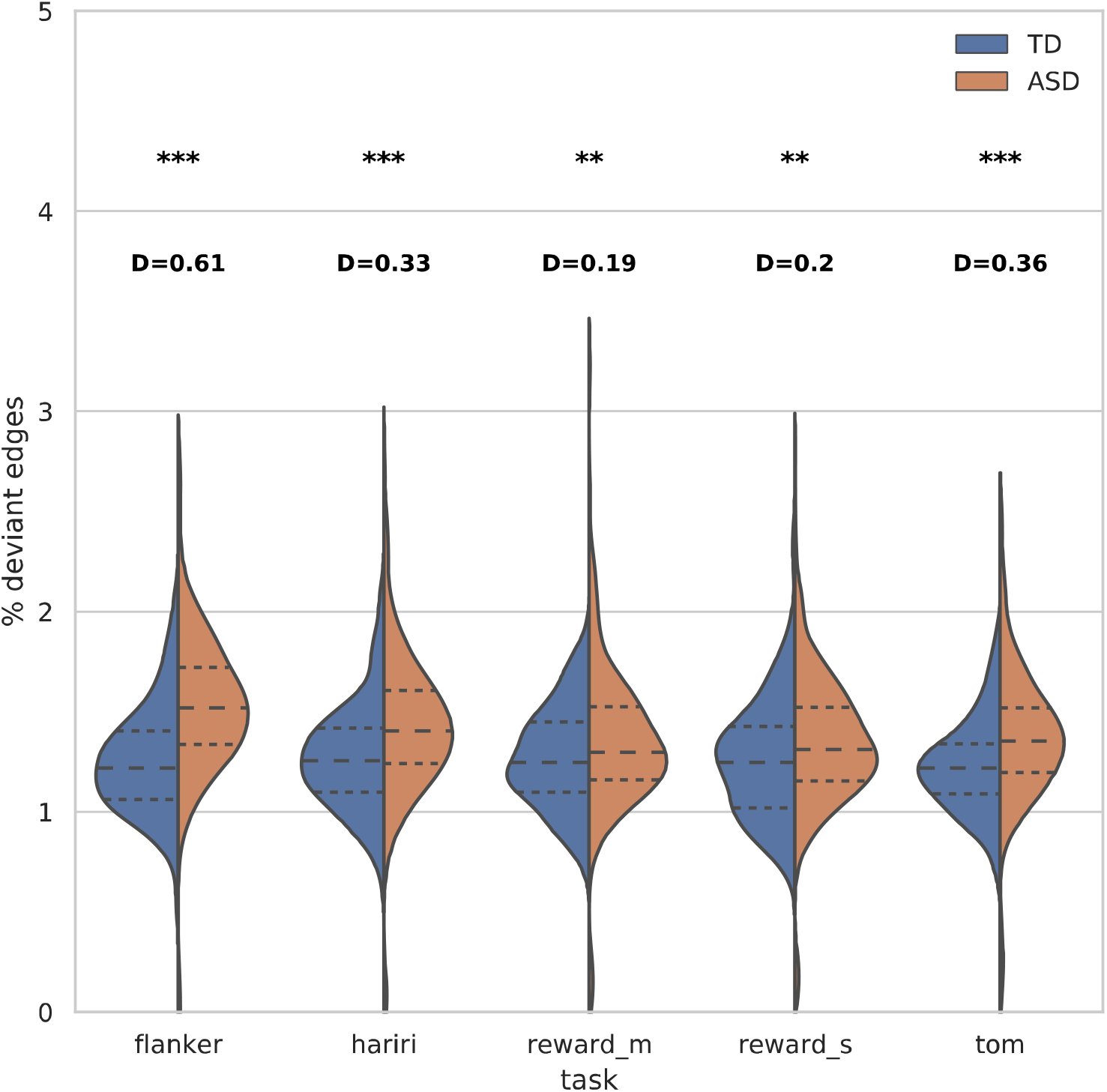
Violin plot of subject distributions for atypicality subject scores in each task. Independent t-tests are performed between the subject distributions for each task. Derived P-values (p), and Cohen’s D effect size (D) are displayed above the respective tasks. (** = p<0.01, ***= p<0.001)

### Brain region atypicality levels

For the next analysis we looked more closely at the spatial profile of connectivity modulation atypicality of autistic individuals with respect to typically developing controls. Figure 2 shows for each task the top 10% brain regions with the greatest atypicality scores in the autism group. To further identify the implicated cognitive terms associated with these regions, we used the Neurosynth online brain image decoder (October, 2021) to identify which networks and cognitive terms were represented in the atypicality data spatial pattern (Gorgolewski et al., 2015). Because atypicality scores were initially estimated at the edge level, we computed the mean absolute atypicality value for all edges in a region, in order to present an atypicality score per brain region. For the Hariri task the top matches were medial prefrontal cortex (r=0.141), auditory (r=0.132), and speech networks (r=0.13), and superior temporal cortex (r=0.119). In the Flanker task, the top matches between the pattern of atypicality and canonical brain networks were in order: medial prefrontal cortex (r=0.144), default mode network (r=0.12), and posterior cingulate (r=0.11). For the monetary reward task, the top matches were again medial prefrontal cortex (r=0.146), then somatosensory cortex (r=0.116), default mode network (r=0.115), and speech networks (r=0.11). For the social reward task, the matches consisted of medial prefrontal cortex (r=0.129), speech network (r=0.11), and default mode network (r=0.108). Finally, in the theory of mind task the spatial pattern of atypical modulation was most closely matched to the speech network (r=0.141), superior temporal cortex (r=0.14), and auditory cortex (r=0.139). The purely cognitive terms associated most with the atypicality scores were speech and listening/auditory regions across all tasks.

**Figure 2.**
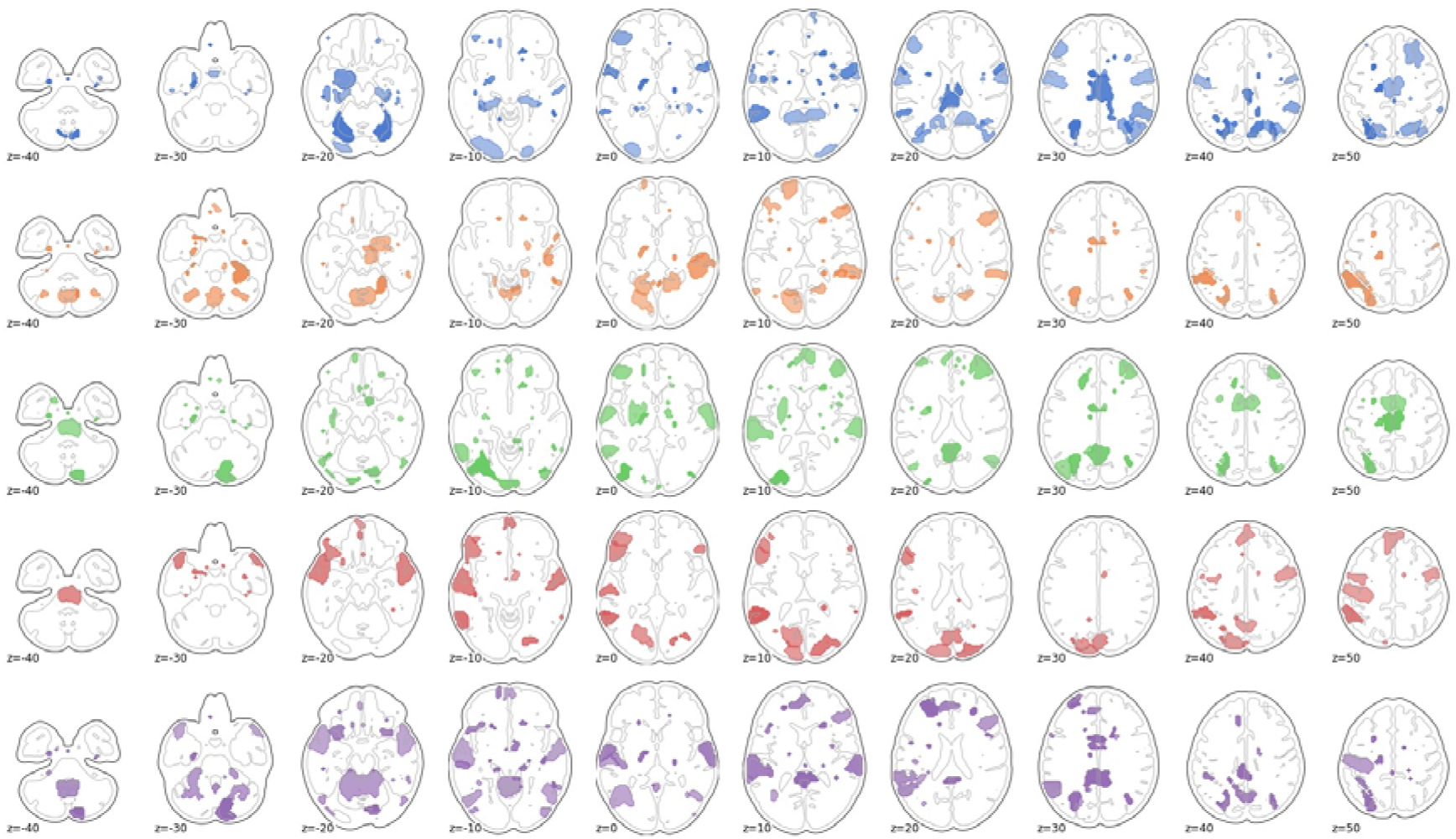
The top 10% brain regions with the greatest atypicality score in autism as summed over edges. From top to bottom: Hariri, Flanker, social reward, nonsocial reward, and theory of mind.

### Atypicality similarity across tasks

Figure 3 shows the correlation matrix of the atypicality edge pattern across the different tasks in the typically developing controls and autism groups. Correlations are consistently across tasks higher for autism (mean correlation: 0.43) than for TD (mean correlation 0.07). Wilcoxon signed rank test of the difference shows a significant difference at p<0.002.

**Figure 3.**
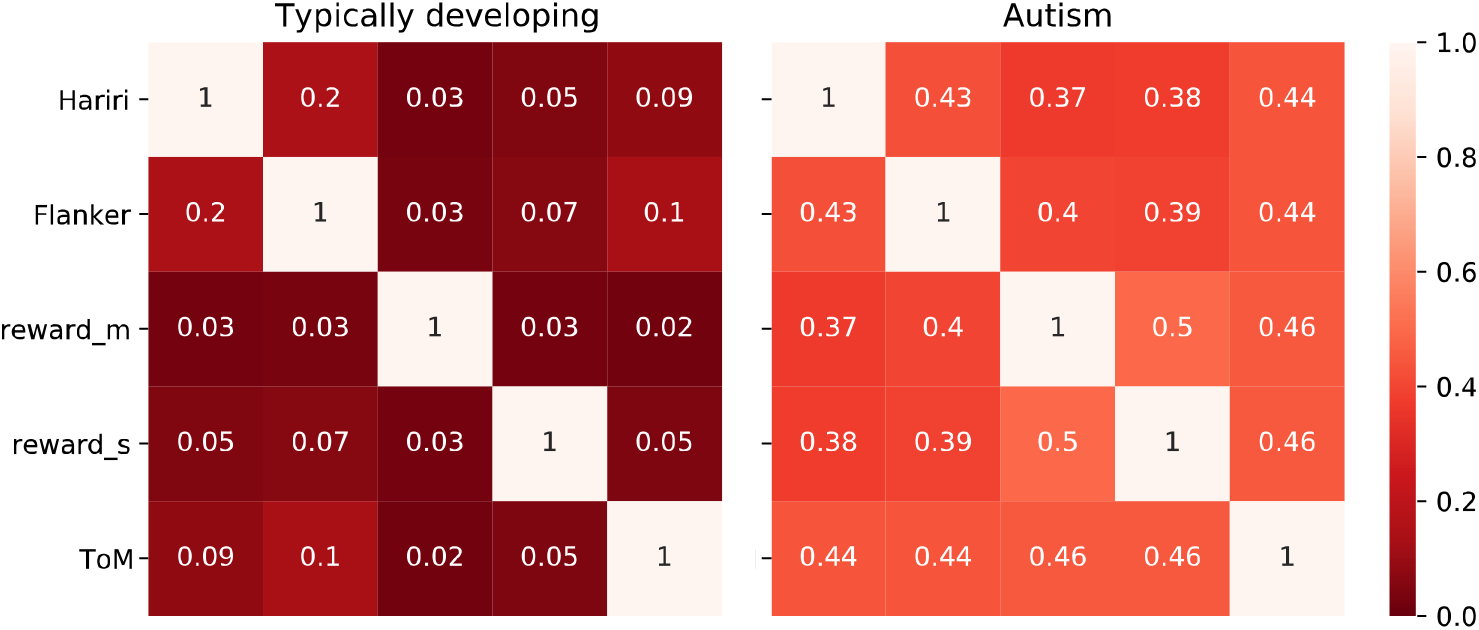
Heatmap of cross-task Pearson correlation of the edgewise group-level mean atypicality pattern in the brain for TD and ASD. Correlations are higher across the board for autism (mean correlation: 0.43), than for TD (mean correlation 0.07). Wilcoxon signed rank test shows a significant difference at p<0.002.

### Canonical correlation analysis

We then investigated whether these spatial patterns of atypicality in the different tasks are also meaningfully related to behavioural measures of autism. This grounds the model into clinical relevance. Canonical correlation analyses found significant brain-behaviour modes of covariation in each of the tasks. Loadings displayed very high stability under out-of-sample 10-fold cross-validation (correlations > 0.99). Figure 4 displays CCA loadings in the behavioural domain for the respective tasks. The pattern found is similar across tasks and follows a trend of autism severity (the SSP is scored inverted relative to the other metrics, with low scores indicating high impairment). The loadings are of comparable magnitude across the tasks with the exception of IQ, which does load in the modes revealed from the Hariri and Flanker tasks, but shows minimal involvement in (non)social reward and theory of mind. Figure 5 shows the top loading brain regions in the CCA. Figure 5 is effectively a rotation of the data from figure 2 under the added influence of autism behavioural scores. In this situation, for the Hariri task the top Neurosynth matches were anterior cingulate cortex (r=0.215), speech cortex (r=0.143), and anterior insula (r=0.134). For the Flanker task these were temporal cortex (r=0.152), language regions (r=0.148), and superior temporal cortex (r=0.148). In the monetary reward task they were visual cortex (r=0.185), occipital cortex (r=0.177), and motor cortex (r=0.162). In social reward the best matches were prefrontal cortex (r=0.135), linguistic regions (r=0.132), and motor cortex (r=0.132). Finally, for the theory of mind task they consisted of dorsolateral cortex (r=0.162), auditory cortex (r=0.145), and anterior cingulate (r=0.14). Supplementary figure 1 shows the difference between the data from figures 2 and 5 visualized.

**Figure 4.**
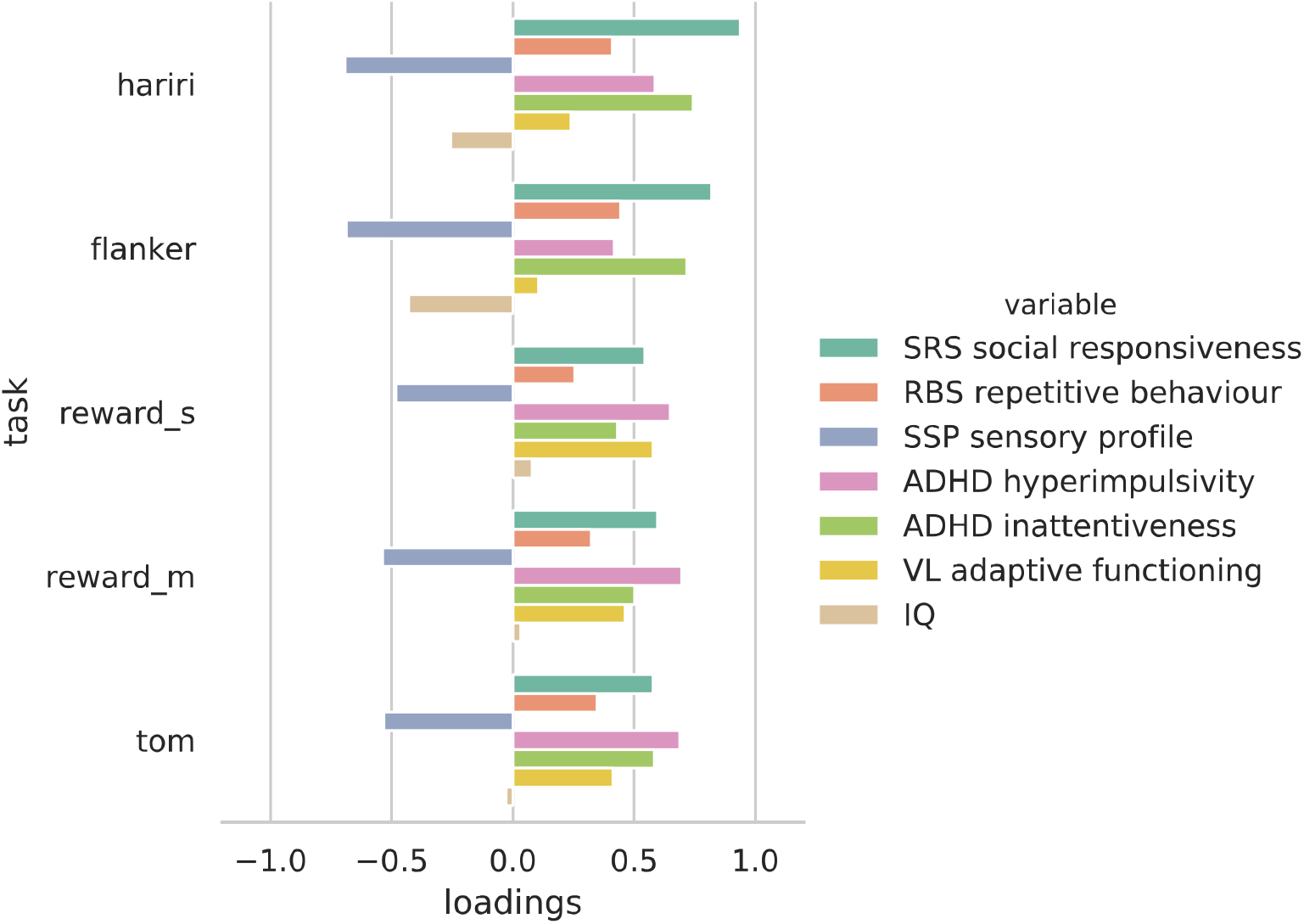
Behavioural loadings for each task in the CCA analysis.

**Figure 5.**
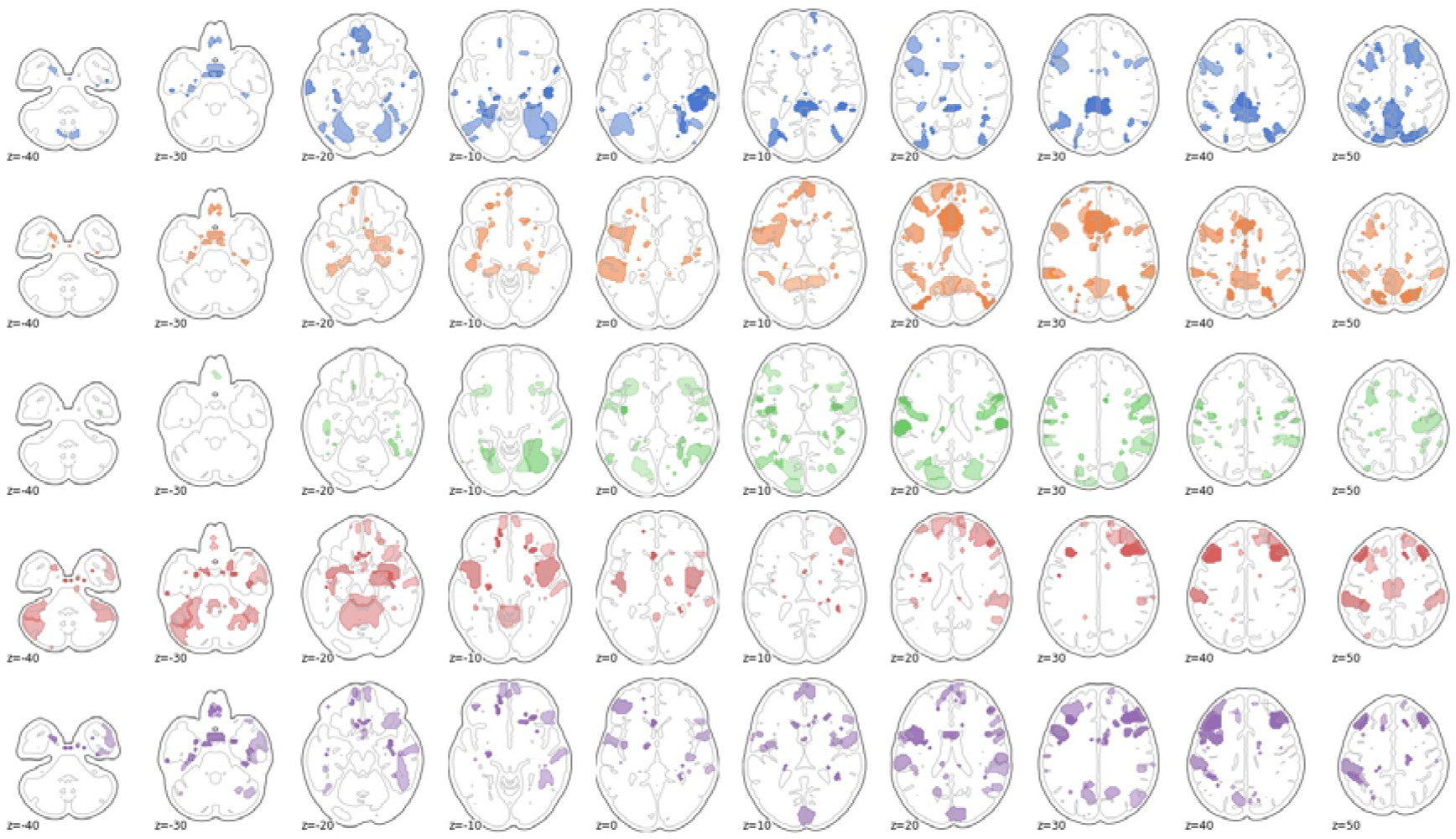
The top 10% brain regions with the greatest CCA loadings in autism as summed over edges. From top to bottom: Hariri, Flanker, social reward, nonsocial reward, and theory of mind.

## Discussion

Our aim in this study was to identify and map individual-level (a)typicality in neural patterns associated with processing across different cognitive domains. We employ a task potency approach (functional connectivity modulation away from resting state baseline) for individuals with autism as they engage in tasks with different cognitive demands.

We show that autism is paired with greater individual-level global atypicality of brain connectivity modulation in each of the tasks under review. This could indicate broadly atypical deployment of neural resources under cognitive loads in autism and reiterates the necessity to move beyond simple group comparisons through the normative modelling approach. We further show that the deviations we model in the brain relate to behavioural measures of autism and found a robust and stable primary relationship in each of the tasks. The behavioural loadings of this relationship can furthermore be interpreted as describing a main axis of impairment in autism.

The spatial patterns of atypicality in the autism group display high levels of cross-task correlation, which was mostly absent in the typically developing controls. This suggests a high level of similar atypicality in autism when dealing with the various cognitive demands. Interestingly, these findings in combination would suggest that while individuals with autism have a globally more atypical pattern of task potency relative to controls, the specific spatial pattern of this atypicality does show clear similarity across cognitive domains. In the context of heterogeneous samples, it lends validity to the concept of an autism as a grouping from a functional neurobiological perspective. Research simultaneously investigating multiple tasks has previously been demonstrated in an ADHD cohort (Chauvin et al., 2021), yet has not been applied to autistic individuals. Given our findings this could be a promising target for future research in autism. A deeper investigation of why and how it is the case that cross-task similarity in atypicalities are on average much more similar in autism than in controls, and how this might relate to literature of the lack of functional differentiation in autism (Nebel et al., 2012; Picci et al., 2016) is necessary. One explanation could be that the brain connectivity pattern of individuals with autism is less free to fluctuate and reorganise under different cognitive loads. This explains both the atypicality in relationship to controls as well as the similarity across tasks within autism as found in this paper.

The findings in this paper need to be contextualised with regards to some limitations. The brain plots displayed in this paper should not be regarded as directly equivalent to activation maps. While areas highlighted in the figures do imply involvement of said area, this involvement is through the up- or downregulation of its coupling with other areas – not necessarily its own activation in isolation. fMRI-tasks used in the LEAP sample were chosen based on their relevance for autism research, however still other forms of cognitive engagement may be relevant for a complete cross-task perspective. Furthermore, though we view autism through individualized atypicality metrics, the normative range estimation necessitates that the typically developing participants are treated as a single group. This potentially masks subgroups in autism that are embedded in the typically developing range.

To conclude, in this paper we have applied innovative techniques to aid understanding of autism brain connectivity heterogeneity in a multi-task setting. These techniques reveal that individuals with autism engage with tasks in a globally atypical way, but that the particular pattern of this atypicality is nevertheless similar across tasks. Atypicalities across tasks originate mostly from prefrontal cortex and default mode network regions, but also speech and auditory networks. We furthermore validated the behavioural relevance of these techniques through showing significant relationships between brain and behavioural data. The similarities between atypicalities across the affected cognitive domains in autism may hold the key to furthering our understanding of the autistic brain. Further, we demonstrated the added value of innovative tools, i.e., task-potency and normative modelling, with the goal to improve the interpretability of task-based fMRI functional connectivity and parse heterogeneity at the individual level in autism. We show that individuals with autism exhibit an atypical task-active functional connectome and we show that taking a cross-task perspective might help reveal a common pattern of atypicality in autism more broadly.

## Acknowledgements

The results leading to this publication have received funding from the Innovative Medicines Initiative 2 Joint Undertaking under grant agreement Nos 115300 (for EU-AIMS) and 777394 (for AIMS-2-TRIALS). This Joint Undertaking receives support from the European Union’s Horizon 2020 research and innovation programme and EFPIA and AUTISM SPEAKS, Autistica, SFARI. DLF is supported by funding from the European Union’s Horizon 2020 research and innovation programme under the Marie Skłodowska-Curie grant agreement No 101025785. This work has been further supported by the European Union Seventh Framework Programme Grant Nos. 602805 (AGGRESSOTYPE) (to JKB), 603016 (MATRICS) (to JKB), and 278948 (TACTICS) (to JKB); European Community’s Horizon 2020 Programme (H2020/2014-2020) Grant Nos. 643051 (MiND) (to JKB), 642996 (BRAINVIEW) (to JKB) and 847818 (CANDY) (to JKB and CFB); the Netherlands Organization for Scientific Research VICI Grant No. 17854 (to CFB); Wellcome Trust Collaborative Award Grant No. 215573/Z/19/Z (to CFB); the Autism Research Trust (to SBC), and the NWO Gravitation Programme Language in Interaction (grant 024.001.006). We thank all participants and their families for participating in this study. We gratefully acknowledge the contributions of all members of the EU-AIMS LEAP group: Jumana Ahmad, Sara Ambrosino, Bonnie Auyeung, Tobias Banaschewski, Simon Baron-Cohen, Sarah Baumeister, Christian F. Beckmann, Sven Bölte, Thomas Bourgeron, Carsten Bours, Michael Brammer, Daniel Brandeis, Claudia Brogna, Yvette de Bruijn, Jan K. Buitelaar, Bhismadev Chakrabarti, Tony Charman, Ineke Cornelissen, Daisy Crawley, Flavio Dell’Acqua, Guillaume Dumas, Sarah Durston, Christine Ecker, Jessica Faulkner, Vincent Frouin, Pilar Garcés, David Goyard, Lindsay Ham, Hannah Hayward, Joerg Hipp, Rosemary Holt, Mark H. Johnson, Emily J.H. Jones, Prantik Kundu, Meng-Chuan Lai, Xavier Liogier D’ardhuy, Michael V. Lombardo, Eva Loth, David J. Lythgoe, René Mandl, Andre Marquand, Luke Mason, Maarten Mennes, Andreas Meyer-Lindenberg, Carolin Moessnang, Nico Mueller, Declan G.M. Murphy, Bethany Oakley, Laurence O’Dwyer, Marianne Oldehinkel, Bob Oranje, Gahan Pandina, Antonio M. Persico, Annika Rausch, Barbara Ruggeri, Amber Ruigrok, Jessica Sabet, Roberto Sacco, Antonia San José Cáceres, Emily Simonoff, Will Spooren, Julian Tillmann, Roberto Toro, Heike Tost, Jack Waldman, Steve C.R. Williams, Caroline Wooldridge, Iva Ilioska, Ting Mei and Marcel P. Zwiers. The funders had no role in the design of the study; in the collection, analyses, or interpretation of data; in the writing of the manuscript, or in the decision to publish the results.) Any views expressed are those of the author(s) and not necessarily those of the funders.

## Supplementary information

### fMRI scanning Parameters

Structural images were obtained using a 5.5-minute MPRAGE sequence (TR=2300ms, TE=2.93ms, T1=900ms, voxels size=1.1×1.1×1.2mm, flip angle=9°, matrix size=256×256, FOV=270mm, 176 slices). The rsfMRI scan was acquired using a multi-echo planar imaging (ME-EPI) sequence; TR=2300ms, TE 12ms, 31ms, and 48ms (slight variations are present across centres), flip angle=80°, matrix size=64×64, in-plane resolution=3.8mm, FOV=240mm, 33 axial slices, slice thickness/gap=3.8mm/0.4mm, volumes=200 (UMCU), 215 (KCL, CIMH), or 265 (RUNMC, UCAM).

### fMRI task brief descriptions

Hariri: participants view a trio of faces and are asked to select one of two test faces that expresses the same emotion as the target face.

Flanker: participants press a button corresponding to the direction of an arrow target. The target is flanked by stimuli that point either in the same (congruent) or opposite (incongruent) direction, or by neutral flankers.

Social reward: An arrow cue indicates to the participants whether a given trial provides an opportunity for a win. Participants are then asked to make a fast button press in response to a target. Succesful eligible trials are rewarded with a happy face stimulus.

Social reward: An arrow cue indicates to the participants whether a given trial provides an opportunity for a win. Participants are then asked to make a fast button press in response to a target. Succesful eligible trials are rewarded with a monetary stimulus.

Theory of mind: participants are presented with short videos in which two triangles either move randomly, in a goal-directed manner, or in a goal directed manner that involves that manipulation of thoughts and feelings of the other triangle. Participants are asked to categorize the videos with one of these three labels.

### Subsample characterizations per task

**Table S1.** (hariri)

| variable | ASD mean(std) [min-max] | TD mean(std) [min-max] | post hoc |
| --- | --- | --- | --- |
| age | 18.22(5.74) [7.58-30.0] | 17.55(5.49) [6.89-30.0] | ns |
| IQ | 106.93(16.29) [63.18-148.0] | 108.03(11.99) [76.82-134.0] | ns |
| SRS social responsiveness | 68.15(11.87) [43.0-95.0] | 54.75(10.89) [37.0-67.0] | ASD>TD |
| RBS repetitive behaviour | 13.83(13.21) [0.0-90.0] | 3.6(3.87) [0.0-23.0] | ASD>TD |
| SSP sensory profile | 146.28(24.03) [53.0-190.0] | 166.04(15.55) [138.0-190.0] | ASD<TD |
| ADHD hyperimpulsivity | 2.15(2.64) [0.0-9.0] | 0.52(1.05) [0.0-6.0] | ASD>TD |
| ADHD inattentiveness | 3.79(2.86) [0.0-9.0] | 1.86(1.69) [0.0-7.0] | ASD>TD |
| VL adaptive functioning | 73.19(12.25) [20.0-117.0] | 79.39(11.73) [73.0-126.0] | ASD<TD |

**Table S2.** (flanker)

| variable | ASD mean(std) [min-max] | TD mean(std) [min-max] | post hoc |
| --- | --- | --- | --- |
| age | 19.16(4.77) [12.07-30.0] | 18.75(4.61) [11.18-30.0] | ns |
| IQ | 104.86(16.54) [63.18-148.0] | 108.02(11.58) [76.82-133.0] | ns |
| SRS social responsiveness | 68.19(11.7) [43.0-95.0] | 55.6(10.83) [37.0-67.0] | ASD>TD |
| RBS repetitive behaviour | 14.18(13.43) [0.0-90.0] | 3.78(3.81) [0.0-23.0] | ASD>TD |
| SSP sensory profile | 147.78(23.29) [53.0-190.0] | 165.44(15.63) [146.0-190.0] | ASD<TD |
| ADHD hyperimpulsivity | 2.0(2.42) [0.0-9.0] | 0.5(0.96) [0.0-6.0] | ASD>TD |
| ADHD inattentiveness | 3.81(3.03) [0.0-9.0] | 2.07(1.68) [0.0-7.0] | ASD>TD |
| VL adaptive functioning | 71.67(11.46) [20.0-101.0] | 75.06(3.84) [73.0-114.0] | ASD<TD |

**Table S3.** (social reward)

| variable | ASD mean(std) [min-max] | TD mean(std) [min-max] | post hoc |
| --- | --- | --- | --- |
| age | 16.95(5.51) [7.48-30.0] | 16.71(5.61) [6.89-30.0] | ns |
| IQ | 101.06(18.62) [59.0-148.0] | 106.8(13.82) [70.41-142.0] | ASD<TD |
| SRS social responsiveness | 69.54(11.43) [43.0-95.0] | 54.76(10.84) [37.0-73.0] | ASD>TD |
| RBS repetitive behaviour | 15.27(13.48) [0.0-90.0] | 3.47(3.87) [0.0-23.0] | ASD>TD |
| SSP sensory profile | 143.88(23.1) [53.0-190.0] | 166.47(15.26) [138.0-190.0] | ASD<TD |
| ADHD hyperimpulsivity | 2.38(2.59) [0.0-9.0] | 0.59(1.23) [0.0-7.0] | ASD>TD |
| ADHD inattentiveness | 4.16(3.0) [0.0-9.0] | 1.88(1.71) [0.0-7.0] | ASD>TD |
| VL adaptive functioning | 72.62(12.11) [20.0-117.0] | 81.58(13.69) [66.0-127.0] | ASD<TD |

**Table S4.** (monetary reward)

| variable | ASD mean(std) [min-max] | TD mean(std) [min-max] | post hoc |
| --- | --- | --- | --- |
| age | 16.99(5.49) [7.48-30.0] | 16.71(5.7) [6.89-30.0] | ns |
| IQ | 100.9(18.56) [59.0-148.0] | 106.92(13.95) [70.41-142.0] | ASD<TD |
| SRS social responsiveness | 69.59(11.4) [43.0-95.0] | 54.75(10.74) [37.0-71.0] | ASD>TD |
| RBS repetitive behaviour | 15.14(13.36) [0.0-90.0] | 3.51(3.86) [0.0-23.0] | ASD>TD |
| SSP sensory profile | 144.18(22.73) [53.0-190.0] | 166.28(15.18) [138.0-190.0] | ASD<TD |
| ADHD hyperimpulsivity | 2.38(2.59) [0.0-9.0] | 0.54(1.17) [0.0-7.0] | ASD>TD |
| ADHD inattentiveness | 4.13(2.97) [0.0-9.0] | 1.81(1.64) [0.0-6.0] | ASD>TD |
| VL adaptive functioning | 72.67(12.03) [20.0-117.0] | 81.72(13.77) [66.0-127.0] | ASD<TD |

**Table S5.** (theory of mind)

| variable | ASD mean(std) [min-max] | TD mean(std) [min-max] | post hoc |
| --- | --- | --- | --- |
| age | 16.97(5.75) [7.48-30.0] | 16.64(5.47) [6.89-30.0] | ns |
| IQ | 105.04(16.37) [63.18-148.0] | 108.22(12.11) [74.0-134.0] | ASD<TD |
| SRS social responsiveness | 69.33(11.57) [43.0-95.0] | 54.31(10.87) [37.0-73.0] | ASD>TD |
| RBS repetitive behaviour | 14.65(12.7) [0.0-90.0] | 3.33(3.82) [0.0-23.0] | ASD>TD |
| SSP sensory profile | 144.76(23.56) [53.0-190.0] | 166.94(15.41) [138.0-190.0] | ASD<TD |
| ADHD hyperimpulsivity | 2.41(2.67) [0.0-9.0] | 0.51(1.05) [0.0-6.0] | ASD>TD |
| ADHD inattentiveness | 4.11(2.98) [0.0-9.0] | 1.77(1.67) [0.0-7.0] | ASD>TD |
| VL adaptive functioning | 73.69(11.67) [20.0-117.0] | 81.29(13.05) [73.0-126.0] | ASD<TD |

### Full statistical values for figure 1

**Table T1.**
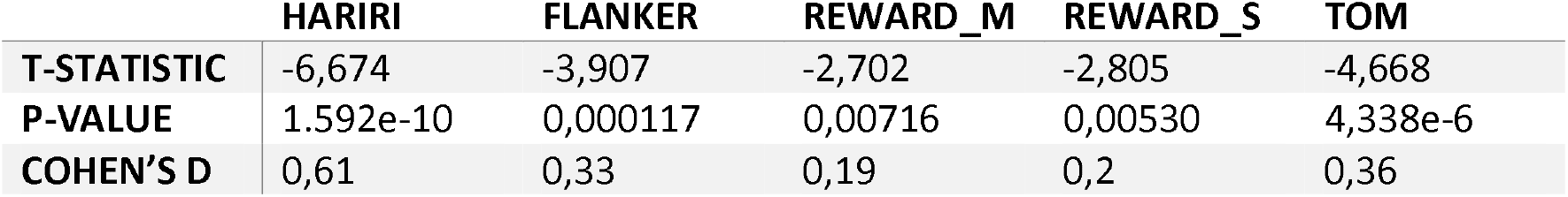

|  | HARIRI | FLANKER | REWARD_M | REWARD_S | TOM |
| --- | --- | --- | --- | --- | --- |
| **T-STATISTIC** | -6,674 | -3,907 | -2,702 | -2,805 | -4,668 |
| **P-VALUE** | 1.592e-10 | 0,000117 | 0,00716 | 0,00530 | 4,338e-6 |
| **COHEN'S D** | 0,61 | 0,33 | 0,19 | 0,2 | 0,36 |

### Supplementary figures

**Figure supp 1.**
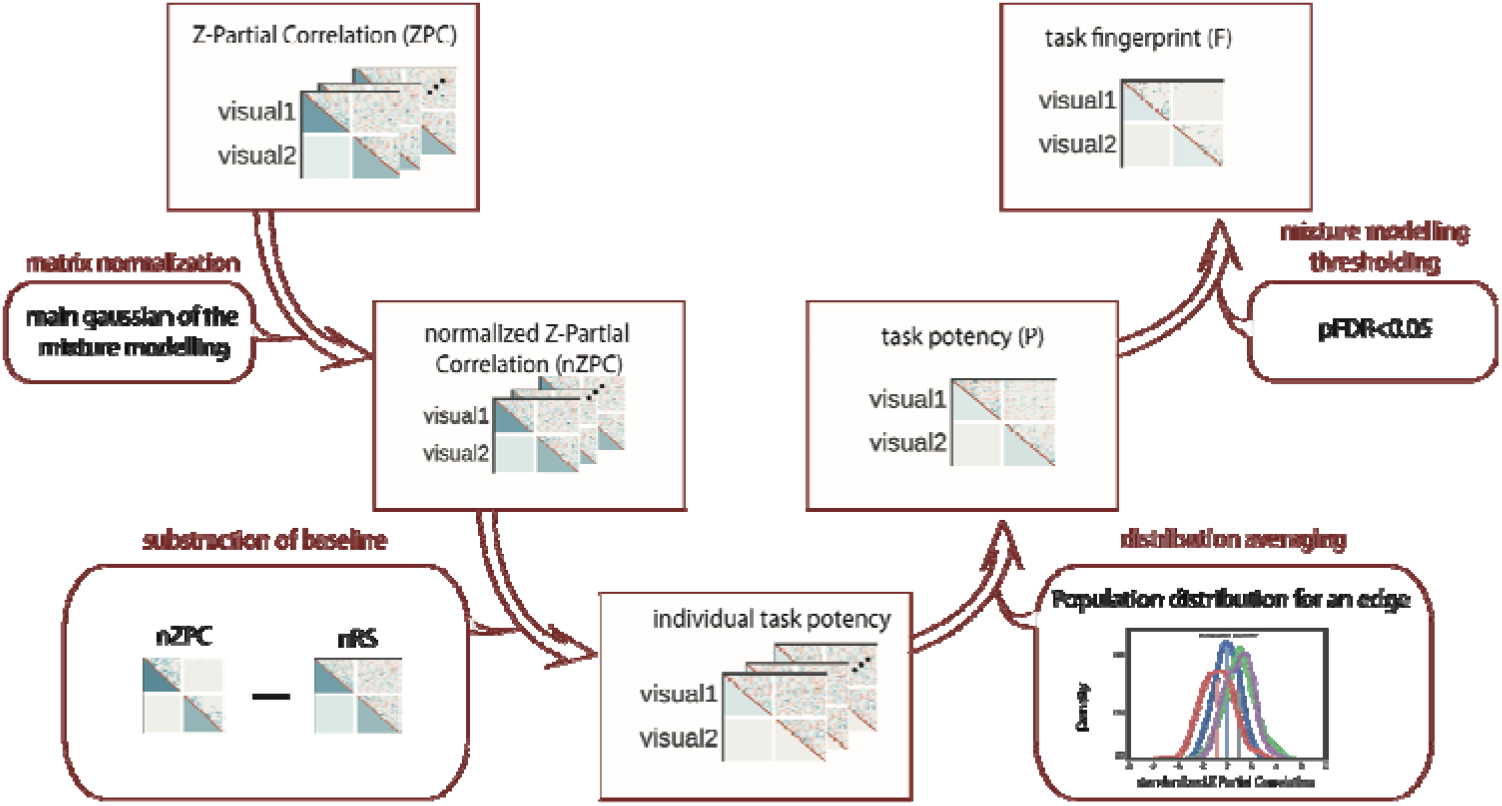
Visual analytical pipeline for task-potency

**Figure supp 2.**
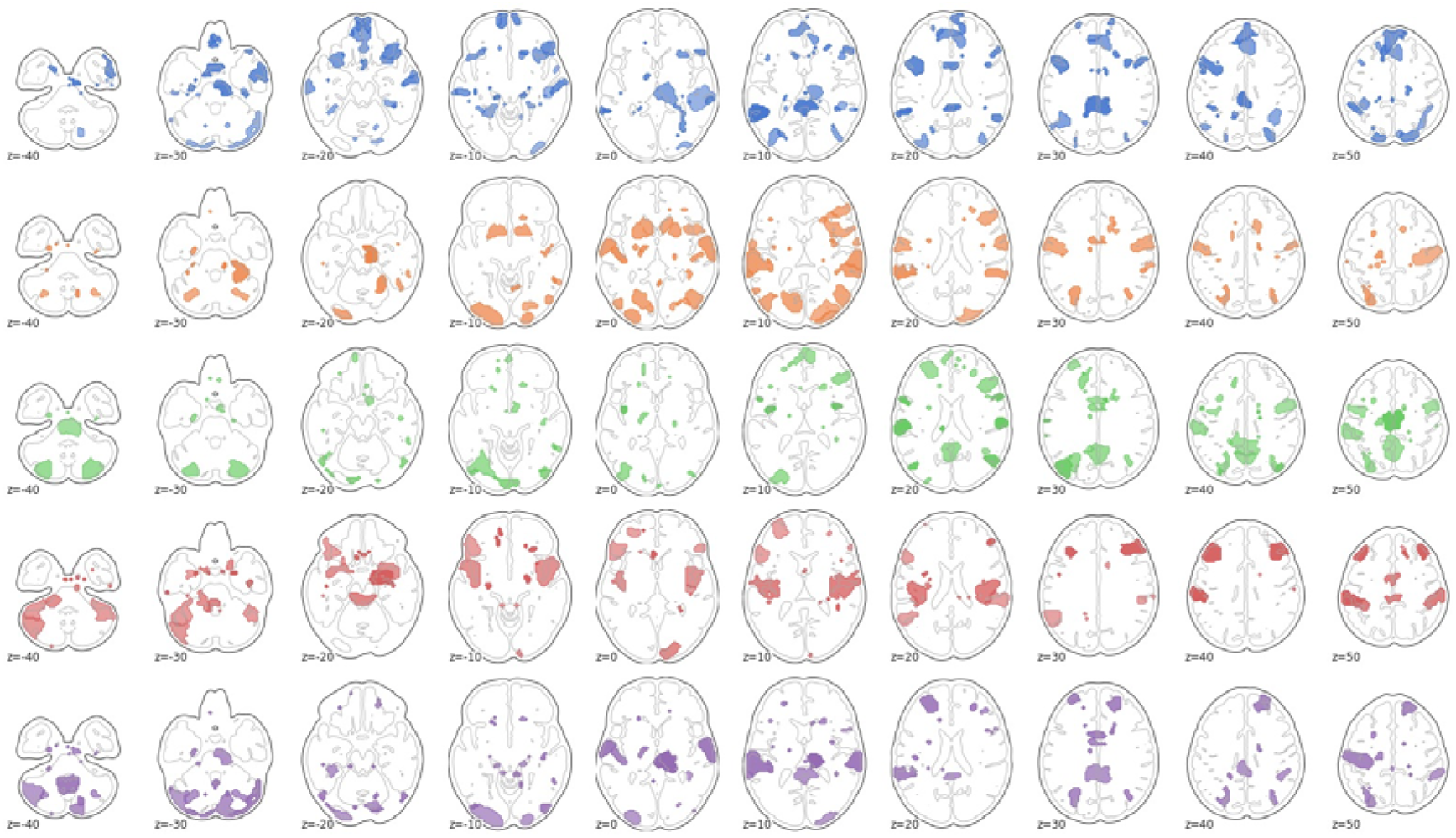
The top 10% brain regions with the greatest difference between normative modeling atypicality scores and CCA loadings in autism as summed over edges. From top to bottom: Hariri, Flanker, social reward, nonsocial reward, and theory of mind.

